# Mind-wandering in people with hippocampal damage

**DOI:** 10.1101/159681

**Authors:** Cornelia McCormick, Clive R. Rosenthal, Thomas D. Miller, Eleanor A. Maguire

## Abstract

Subjective inner experiences, such as mind-wandering, represent the fundaments of human cognition. Although the precise function of mind-wandering is still debated, it is increasingly acknowledged to have influence across cognition on processes such as future planning, creative thinking and problem-solving, and even on depressive rumination and other mental health disorders. Recently, there has been important progress in characterizing mind-wandering and identifying the associated neural networks. Two prominent features of mind-wandering are mental time travel and visuo-spatial imagery, which are often linked with the hippocampus. People with selective bilateral hippocampal damage cannot vividly recall events from their past, envision their future or imagine fictitious scenes. This raises the question of whether the hippocampus plays a causal role in mind-wandering and if so, in what way. Leveraging a unique opportunity to shadow people (all males) with bilateral hippocampal damage for several days, we examined, for the first time, what they thought about spontaneously, without direct task demands. We found that they engaged in as much mind-wandering as control participants. However, whereas controls thought about the past, present and future, imagining vivid visual scenes, hippocampal damage resulted in thoughts primarily about the present comprising verbally-mediated semantic knowledge. These findings expose the hippocampus as a key pillar in the neural architecture of mind-wandering and also reveal its impact beyond memory, placing it at the heart of human mental life.

**Significance statement:** Humans tend to mind-wander about 30-50% of their waking time. Two prominent features of this pervasive form of thought are mental time travel and visuo-spatial imagery, which are often associated with the hippocampus. To examine whether the hippocampus plays a causal role in mind-wandering, we examined the frequency and phenomenology of mind-wandering in patients with selective bilateral hippocampal damage. We found that they engaged in as much mind-wandering as controls. However, hippocampal damage changed the form and content of mind-wandering from flexible, episodic, and scene-based to abstract, semanticized, and verbal. These findings expose the hippocampus as a key pillar in the neural architecture of mind-wandering and reveal its impact beyond memory, placing it at the heart of our mental life.

## Introduction

Even when in the same place and involved in the same activity, at any given moment people can experience the world in different ways. Recently, there have been advances in delineating the various forms of spontaneous inner experiences and their neural correlates (Andrews-Hanna et al., 2014a; Christoff et al., 2016). Self-generated thinking typically refers to the ability to mentally decouple from current perceptual surroundings and generate independent internal thoughts (Smallwood and Schooler, 2015). These thoughts can either be task-related, such as actively thinking about how this manuscript should be structured, or task-unrelated, where there is a spontaneous inner focus, such as suddenly remembering what a nice time I had yesterday with my friends (Seli et al., 2016). These latter thoughts are the focus of the current study, and have been variously described as task-unrelated self-generated thoughts, daydreaming or mind-wandering (Smallwood and Schooler, 2015).

It has been shown that humans tend to mind-wander about 30-50% of waking time, irrespective of the current activity (Kane et al., 2007; Killingsworth and Gilbert, 2010). Nevertheless, mind-wandering frequency is particularly pronounced during restful periods and low-demanding tasks (Smallwood and Schooler, 2015). The latter is often exploited by experimentalists examining mind-wandering. Although the precise function of mind-wandering is still debated, it is increasingly acknowledged to have influence across cognition on processes such as future planning, creative thinking and problem-solving (Baird et al., 2011; Baird et al., 2012), and even on depressive rumination and other mental health disorders (Ehlers et al., 2004; Andrews-Hanna et al., 2014a). Furthermore, the content of mind-wandering seems wide-ranging, including episodic memory recall (which involves a sense of re-experiencing and is specific in time and place), future planning, mentalizing, simulation of hypothetical scenarios, and involves a variety of emotions and different sensory modalities (Andrews-Hanna et al., 2013; Smallwood et al., 2016). Interestingly, two of the most prominent features of mind-wandering are mentally travelling forwards and backwards in time and visual imagery, which are functions usually associated with the hippocampus (Tulving, 1985, 2002; Hassabis et al., 2007).

The Default Mode Network (DMN), within which the hippocampus is a node, has been associated with self-generated thoughts such as mind-wandering (Buckner et al., 2008; Andrews-Hanna et al., 2014b). Of particular relevance here, stronger hippocampal connectivity with other regions of the DMN was observed in individuals who experienced more episodic details and greater flexibility in mental time travel during mind-wandering episodes (Karapanagiotidis et al., 2016; Smallwood et al., 2016). Unfortunately, causal evidence for hippocampal involvement in mind-wandering is lacking (Fox et al., 2016). Behavioral studies of patients with lesions are crucial because they permit examination of the causal effects of regional brain damage on the networks established by neuroimaging work. People with hippocampal damage cannot vividly recall events from their past (Lah and Miller, 2008) envision their future (Kurczek et al., 2015) or imagine fictitious scenes (Hassabis et al., 2007). Therefore, whether they experience mind-wandering, and if they do, what form does it take, are important and timely questions which we addressed by examining mind-wandering in patients with selective bilateral hippocampal damage.

Previous studies have examined the effects of hippocampal damage during demanding tasks, such as autobiographical memory retrieval (Lah and Miller, 2008), designed to challenge the patients’ cognitive abilities. In contrast, in order to establish what patients with hippocampal damage think about spontaneously when there is no concurrent task, our focus was on what they do in their mentally “free” time. We initially asked whether or not patients with hippocampal damage were able to mentally decouple from the current perceptual input. If yes, we then had a series of further questions. First, would they engage in mental time travel? Second, what form would their mind-wandering take – spontaneous episodic, detailed thoughts or semantic, abstract thoughts? Lastly, we asked whether they experienced spontaneous visual imagery similar to that typically reported by control participants during mind-wandering (Andrews-Hanna et al., 2013)?

## Materials and Methods

### Participants

Six patients (all right-handed males, mean age 57.0 years (SD 16.9), age range 27 to 70) with selective bilateral hippocampal lesions and selective episodic memory impairment took part (see Tables 1 and 2 for demographic information and neuropsychological profiles). Of note, these patients were the same high-functioning individuals that took part in our previous studies (McCormick et al., 2016, 2017a). Hippocampal damage (see example in Fig. 1a) resulted in all cases from voltage-gated potassium channel (VGKC)-complex antibody-mediated limbic encephalitis (LE). Two of the patients had bilateral signal hyperintensities in the hippocampi on presentation, but hippocampal atrophy was observed in all patients. Testing took place a median of seven years post-onset of hippocampal damage. In line with previous reports of this patient population (Dalmau and Rosenfeld, 2014; Miller et al., 2017), manual (blinded) segmentation of the hippocampi from high-resolution structural MRI scans confirmed that our patients showed volume loss confined to the left (Patients – HPC: 2506mm^3^ (mean) +/-394 (standard deviation), control participants – CTL: 3173 mm^3^ +/-339, W=4.0, p=0.002) and right (HPC: 2678mm^3^ +/-528, CTL: 3286mm^3^ +/-301, W=8.0, p=0.01) hippocampus. To rule out pathological differences between patients and controls elsewhere in the brain, an automated voxel-based-morphometry (VBM; Ashburner, 2009) analysis was carried out on whole brain T1 weighted MRI images and, in line with previous reports on patients of this sort (Wagner et al., 2015; Finke et al., 2017; Miller et al., 2017), did not result in any significant group differences outside of the hippocampus even at a liberal uncorrected p-value of less than 0.001.

**Table 1.** Summary of demographic information.

| **Group** | **N** | **HD** | **Age** | **Chronicity** | **LHPC vol\*** | **RHPC vol\*** |
| --- | --- | --- | --- | --- | --- | --- |
| HPC group | 6 (M) | 6 (R) | 57.0<br>(16.9) | 6.8<br>(2.1) | **2506**<br>(394) | **2678**<br>(528) |
| CTL group | 12 (M) | 11 (R) | 57.2<br>(16.6) | n.a. | **3173**<br>(339) | **3286**<br>(301) |
| p-value |  |  | 0.97 | n.a. | **0.002** | **0.01** |

For both groups, means are displayed with standard deviations below the corresponding mean in parentheses. HPC=hippocampal-damaged patients; CTL=healthy control participants; M=Male; HD=Handedness; n.a.=not applicable; R=Right; L=Left; vol=volume in mm<sup>3</sup>. \*One control participant could not be scanned, therefore hippocampal volumes are based on all six patients and 11 control participants. Age and chronicity are described in years. p-value=p-value of between-group non-parametric Mann-Whitney U tests with significant differences depicted in bold.

**Table 2.** Summary of neuropsychological information.

|  | Controls |  | HPC Patients |  | P-Value |
| --- | --- | --- | --- | --- | --- |
|  | M | SD | M | SD |  |
| General Cognition |  |  |  |  |  |
| WASI Matrix Reasoning | 13.8 | 1.5 | 13.2 | 2.2 | 0.51 |
| WASI Similarities | 11.8 | 2.6 | 12.8 | 1.8 | 0.54 |
| Episodic memory |  |  |  |  |  |
| Autobiographical Interview int* | 51.3 | 13.6 | 31.7 | 6.7 | 0.01 |
| Autobiographical Interview ext* | 5.9 | 2.2 | 6.1 | 3.8 | 0.67 |
| WMS Logical Memory (immediate recall, units) | 12.6 | 3.2 | 8.7 | 2.4 | 0.03 |
| WMS Logical Memory (immediate recall, thematic) | 13.8 | 3.0 | 9.2 | 2.6 | 0.01 |
| WMS Wordlist (immediate recall) | 13.3 | 3.2 | 10.2 | 3.9 | 0.14 |
| Rey-Osterrieth Complex Figure (copy /36) | 35.5 | 1.4 | 33.4 | 4.2 | 0.19 |
| Rey-Osterrieth Complex Figure (immediate recall /36) | 23.8 | 7.4 | 19.0 | 5.9 | 0.12 |
| WMS Logical Memory (delayed recall, units) | 13.2 | 3.7 | 7.8 | 4.0 | 0.01 |
| WMS Logical Memory (delayed recall, thematic) | 13.5 | 3.2 | 7.0 | 4.8 | 0.01 |
| WMS Wordlist (delayed recognition) | 11.7 | 1.4 | 9.3 | 4.5 | 0.47 |
| Rey-Osterrieth Complex Figure (delayed recall, /36) | 23.8 | 7.8 | 18.1 | 6.4 | 0.06 |
| Warrington Recognition Memory test for Words | 12.0 | 2.3 | 12.3 | 2.4 | 0.99 |
| Warrington Recognition Memory test for Faces | 11.3 | 3.0 | 9.2 | 4.3 | 0.14 |
| Semantic memory |  |  |  |  |  |
| Warrington Graded Naming Test | 13.7 | 2.4 | 12.5 | 3.0 | 0.34 |
| Attention/Working memory |  |  |  |  |  |
| WMS Digit Span (forward) | 13.3 | 3.2 | 12.0 | 2.5 | 0.76 |
| Executive Functions |  |  |  |  |  |
| DKEFS Letter Fluency (FAS) | 14.3 | 3.2 | 12.7 | 3.7 | 0.32 |
| DKEFS Category Fluency | 13.9 | 4.4 | 12.5 | 5.2 | 0.51 |
| DKEFS Category Switch Test | 12.7 | 2.9 | 12.3 | 3.5 | 0.59 |
| DKEFS Stroop Word-Colour Interference Test | 12.4 | 2.2 | 13.3 | 2.2 | 0.41 |
| Hayling Sentence Completion Test (coherent) | 6.1 | 1.0 | 5.8 | 0.4 | 0.73 |
| Hayling Sentence Completion Test (incoherent) | 5.8 | 0.8 | 5.8 | 0.4 | 0.95 |
| Hayling Sentence Completion Test (errors) | 6.8 | 1.1 | 6.5 | 1.9 | 0.93 |
| Hayling Sentence Completion Test (total) | 18.7 | 1.6 | 18.2 | 2.1 | 0.75 |
| DKEFS Trails Test (visual scanning) | 12.0 | 1.3 | 11.2 | 1.0 | 0.17 |
| DKEFS Trails Test (number sequencing) | 11.8 | 2.5 | 10.2 | 2.3 | 0.12 |
| DKEFS Trails Test (letter sequencing) | 12.5 | 1.5 | 11.0 | 2.4 | 0.21 |
| DKEFS Trails Test (letter-number sequencing) | 12.8 | 1.0 | 10.7 | 2.2 | **0.04** |
| DKEFS Trails Test (motor speed) | 11.8 | 1.1 | 10.2 | 4.5 | 0.99 |
| **Visual perception** |  |  |  |  |  |
| VOSP Dot Counting (/10) | 10.0 | 0.0 | 9.7 | 0.8 | 0.33 |
| VOSP Position Discrimination (/20) | 20.0 | 0.0 | 19.7 | 0.8 | 0.33 |
| VOSP Cube Analysis (/10) | 9.6 | 0.8 | 9.7 | 0.8 | 0.99 |
| VOSP Overall (/40) | 39.6 | 0.8 | 39.0 | 2.4 | 0.99 |
| **Mood** |  |  |  |  |  |
| HADS Anxiety | 4.3 | 2.9 | 4.3 | 3.3 | 0.83 |
| HADS Depression | 2.3 | 2.7 | 2.5 | 2.3 | 0.71 |

**Figure 1.**
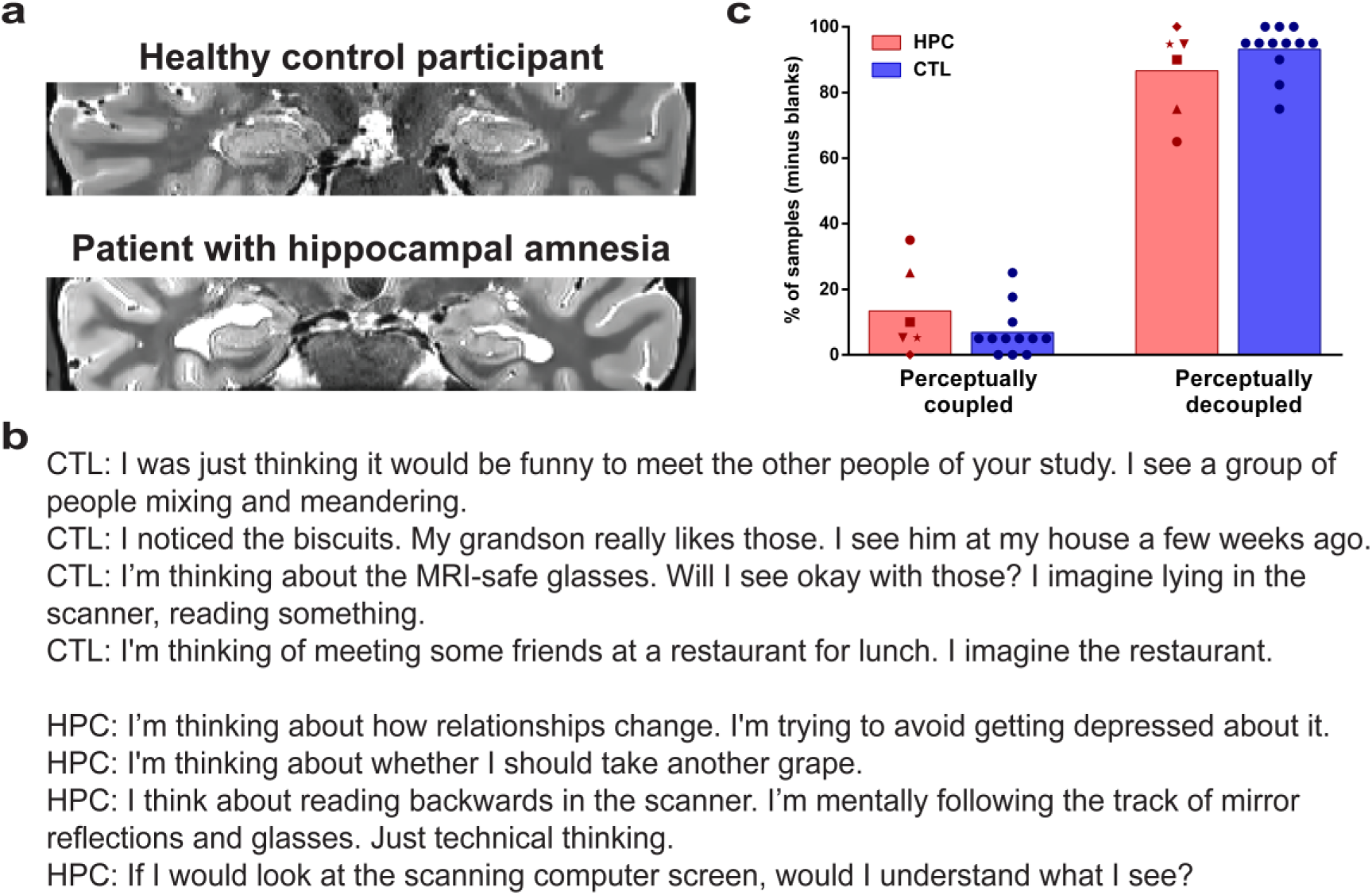
Hippocampal damage and the frequency of mind-wandering. **(a)** A T2-weighted structural MR image of an example patient with selective bilateral hippocampal damage and an age, gender and IQ-matched healthy control participant. Images are displayed in native space corresponding approximately to the position of y=-10 in the MNI coordinate system. **(b)** Examples of mind-wandering experiences from controls (CTL) and patients with hippocampal damage (HPC). **(c)** The average percentage of perceptually coupled and decoupled spontaneous thoughts (minus ‘blank’ thoughts) during quiet restful moments for individual patients with hippocampal damage (red symbols) and healthy control participants (blue circles). Both groups reported a high level of mind-wandering experiences, with no differences between patients and control participants.

Neuropsychologically, the patients displayed an impairment in immediate and delayed recall on the Logical Memory (short stories) test (Wechsler, 1997), and they recollected significantly fewer episodic (‘internal’), but not semantic (‘external’) details on the Autobiographical Interview (Levine et al., 2002), as detailed in Table 2. All other cognitive and emotional aspects of cognition were intact in these patients. In summary, these patients seemed to have a selective difficulty in re-constructing internal events. Importantly, their working memory capacity did not differ from that of controls, suggesting that the differences in mind-wandering episodes we report here are unlikely to be due to an inability to remember the thoughts.

Twelve healthy control participants also took part (all male, one left-handed, mean age 57.2 (16.6) years, age range from 25 to 77). In addition to comparing the two groups, we ensured that each patient was matched closely to two of the control subjects on sex, age, and general cognitive ability (measured by the Matrix Reasoning and Similarities subtests of the Wechsler Abbreviated Scale of Intelligence – WASI; Wechsler, 1999). There were no significant differences between patients and controls on age, general cognitive ability and on neuropsychological tests assessing semantic memory, language, perception, executive functions and mood (see Table 2). All participants gave informed written consent in accordance with the local research ethics committees.

### Characterization of hippocampal damage

#### High resolution T2-weighted structural MRI scans of the medial temporal lobes

Five of the patients and 10 of the control participants underwent structural MR imaging limited to a partial volume focused on the temporal lobes using a 3.0-T whole body MR scanner (Magnetom TIM Trio, Siemens Healthcare, Erlangen, Germany) operated with a radiofrequency (RF) transmit body coil and 32-channel head RF receive coil. These structural images were collected using a single-slab 3D T2-weighted turbo spin echo sequence with variable flip angles (SPACE; Mugler et al., 2000) in combination with parallel imaging, to simultaneously achieve a high image resolution of ~500μm, high sampling efficiency and short scan time while maintaining a sufficient signal-to-noise ratio (SNR). After excitation of a single axial slab the image was read out with the following parameters: resolution=0.52 x 0.52 x 0.5 mm, matrix=384 x 328, partitions=104, partition thickness=0.5 mm, partition oversampling=15.4%, field of view=200 x 171 mm 2, TE=353 ms, TR=3200 ms, GRAPPA x 2 in phase-encoding (PE) direction, bandwidth=434 Hz/pixel, echo spacing=4.98 ms, turbo factor in PE direction=177, echo train duration=881, averages=1.9. For reduction of signal bias due to, for example, spatial variation in coil sensitivity profiles, the images were normalized using a prescan, and a weak intensity filter was applied as implemented by the scanner’s manufacturer. It took 12 minutes to obtain a scan.

#### High resolution T1-weighted structural MRI scans of the whole brain at 3.0 Tesla

In addition, five of the patients and 11 of the control participants underwent a whole brain structural T1weighted sequence at an isotropic resolution of 800μm (Callaghan et al., 2015) which was used for the automated VBM analysis (one control participant could not be scanned). These images had a FoV of 256mm head-foot, 224mm anterior-posterior (AP), and 166mm right-left (RL). This sequence was a spoiled multi-echo 3D fast low angle shot (FLASH) acquisition with a flip angle of 21^0^ and a repetition time (TR) of 25ms. To accelerate the data acquisition, partially parallel imaging using the GRAPPA algorithm was employed in each phase-encoded direction (AP and RL) with forty reference lines and a speed up factor of two. Gradient echoes were acquired with alternating readout polarity at eight equidistant echo times ranging from 2.34 to 18.44ms in steps of 2.30ms using a readout bandwidth of 488Hz/pixel (Helms and Dechent, 2009). The first six echoes were averaged to increase SNR (Helms and Dechent, 2009) producing a T1-weighted image with an effective echo time of 8.3 ms.

#### High resolution T1-weighted MRI scans of the whole brain at 7.0 Tesla

One patient could not be scanned at our Centre due to recent dental implants. We therefore used a whole brain T1-weighted image acquired previously on a 7.0 Tesla MRI scanner - a three-dimensional whole-brain T1-weighted phase sensitive inversion recovery sequence (Mougin et al., 2015) at an isotropic resolution of 600μm, with a tailored inversion pulse for magnetization inversion at ultrahigh field (Hurley et al., 2010), providing inherent bias field correction.

#### Hippocampal segmentation

To improve the SNR of the anatomical images, two or three T2-weighted high resolution scans were acquired for a participant. Images from each participant were co-registered and denoised following the Rician noise estimation (Coupe et al., 2010). The denoised images were averaged and smoothed with a full-width at half maximum kernel of 2x2x2mm. In each case, left and right hippocampi were manually (blindly) segmented and volumes extracted using the ITK Snap software version 3.4.0 (Yushkevich et al., 2006).

#### VBM analysis

An automated VBM analysis was performed using SPM12 (Statistical Parametric Mapping, Wellcome Centre for Human Neuroimaging, London, UK). The averaged T1-weighted images were segmented into grey and white matter probability maps using the unified segmentation approach (Ashburner and Friston, 2005). Inter-subject registration of the tissue classes was performed using Dartel, a nonlinear diffeomorphic algorithm (Ashburner, 2007). The resulting Dartel template and deformations were used to normalize the tissue probability maps to the stereotactic space defined by the Montreal Neurological Institute (MNI) template. For VBM analysis, the normalization procedure included modulating the grey matter tissue probability maps by the Jacobian determinants of the deformation field and smoothing with an isotropic Gaussian smoothing kernel of 8 mm full width at half maximum (FWHM). The normalized grey matter from controls and the patients with hippocampal damage were contrasted using a two sample t-test and thresholded at p<0.001 uncorrected and a cluster extend of 50 voxels.

### Experimental design and procedure

We had the opportunity to shadow the patients with selective bilateral hippocampal damage over two days during day-time hours, and so we adapted for use a well-established method, descriptive experience sampling (DES), in which participants are asked frequently over an extended period of time to describe what was on their minds just before they were aware of being asked (Hurlburt, 1979; Hurlburt and Heavey, 2001; Hurlburt and Akhter, 2006). DES has the advantage that thought probes can extend over a long period of time and the sampling interval can be more extensive than alternative approaches in which a few thought samples are taken while participants perform low-demanding distractor tasks (Smallwood et al., 2002; Smallwood and Schooler, 2015). Furthermore, using DES, participants are encouraged to describe freely what was on their minds, rather than categorizing thoughts into pre-specified classes.

To mitigate any potential difficulties the hippocampal-damaged patients may have had with remembering task instructions over longer time-scales, we made a number of adaptations to the original DES protocol. For example, we changed the type of reminder. The reminder is an important tool as it identifies the precise moment of sampling and happens externally to the participant, meaning that the participant does not have to remember to track their own thoughts (Hurlburt and Stuart, 2014). Usually, DES participants carry a beeper and receive frequent sampling reminders while going about their everyday life (Hurlburt and Akhter, 2006). However, we adapted this sampling method to suit an extended experimental setting over two days in which patients and controls experienced the same structured days (three MRI scans, various cognitive tasks, breaks, lunches, etc). In our case, the experimenter provided the external cue for the participant. Equally important as the reminder is the exact time point of the sample. While previous studies have used a random sampling schedule (Hurlburt, 1979; Smallwood and Schooler, 2015), our main goal was to examine the general ability to perceptually decouple and the content of spontaneous thoughts of these rare patients. We therefore tried to maximize our chances of catching perceptually decoupled thoughts. Hence, we probed 20 times over the course of two structured research days (8 hours each) at pre-specified times in restful moments. To keep the experimental context of the sampling time points as closely matched across participants as possible, thoughts for all participants were probed in the same rooms of our Centre, around the same times of day, and in approximately the same experimental situations. This procedure resulted in schedules whereby some samples were separated by several hours (e.g., during which the participant underwent MRI scanning), and other samples which were relatively close in time (e.g., a few minutes). In addition, in order to ascertain that all participants, especially the patients, could remember time spans long enough to report their thoughts, we asked them to describe two experiments unrelated to the current study shortly after completion. All participants were able to provide accurate accounts of those experiments.

During sampling moments, such as after obtaining consent, and at the beginning of the tea break, the experimenter would allow for a moment of quiet to emerge. That is, the experimenter would fill out some forms or naturally disengage from any conversation. When there was an appropriate time of silence, the experimenter would ask the participant “What were you thinking about just before I asked you?” The participant was encouraged to briefly describe the current thought in one or two sentences. On a prepared note sheet, the participant’s response was written down verbatim. In a follow-up question, the experimenter established whether the thought had been a visual image (if yes, scene or object) or a verbal thought. Then, the experimenter clarified whether that thought had concerned the past, present or future (and if it had been past or future, how far into the past or future). The sampling procedure lasted no longer than approximately one minute to prevent lengthy posthoc elaboration. Lastly, divergent from other DES reports, we opted not to train our participants before the start of the study. Although the training may have provided useful guidance in monitoring one’s own thoughts for the control participants, we felt that patients might not find this as beneficial. Therefore, because the experimenter was present for all samples, none of the participants was required to remember the follow-up questions themselves but were instead cued by the experimenter. Of note, control participants reported equal numbers of decoupled, scene-based thoughts in the first and second half of the samples (first half=5.8 +/-0.9, second half=6.3 +/-1.2, W=11.0, p=0.42), suggesting that there was no significant training effect. A lack of monitoring was further confirmed by the control participants, because they anticipated the sampling probe for only 3 out of a total of 240 sampled thoughts.

Whereas previous research has examined the frequency and content of mind-wandering episodes in healthy participants for features such as goal-orientation and emotional valence (Andrews-Hanna et al., 2013; Andrews-Hanna et al., 2014a; Christoff et al., 2016), we focused here on examining the effect of hippocampal damage on the frequency, time range, representational content and form of mind-wandering, which are key to understanding hippocampal function.

In summary, our adapted sampling protocol permitted us to leverage the naturalistic approach of the typical DES reports that sample over an extended period of time, and allowed participants to report their thoughts freely, while equating the daily activities and the sampling moments of patients and controls participants to maximize our chances of catching perceptually decoupled thoughts in an experimentally rigorous manner.

### Scoring

#### Perceptually coupled or decoupled thoughts and mind-blanking

An episode was considered mind-wandering when the response indicated that the mind was disengaged from the external world (perceptually decoupled; Smallwood and Schooler, 2015). For example, the thought “I see your watch” was considered perceptually coupled, whereas the thought “Time is sometimes slow and sometimes fast” was considered perceptually decoupled. In a few instances, patients and control participants reported thinking about nothing (i.e., mind-blanking; Ward and Wegner, 2013). The frequency of this mind-blanking did not differ between the groups (CTL mean 0.4 +/- 0.88, HPC=1.8 +/-2.8, MWU=23, p=0.19), and we therefore excluded these samples from further analysis.

#### Temporal range

After each sample, we clarified directly with participants whether that thought had concerned the past, present or future, and if past or future, how distant into the past or future. We further sorted participants’ responses from the “present” category based on the observation that patients and control participants reported very different types of thoughts. Consequently, we classified each mind-wandering episode that was labelled by participants as concerning the present moment as either an atemporal scenario or not, in line with the protocol of Jackson et al. (2013). A mind-wandering episode was considered “present”-related if the thought was perceptually decoupled but concerned the now, for example “I’m thinking that you are right-handed” or “I wonder whether I should eat another grape”. On the other hand, a thought was classified as an atemporal scenario if the participant reported a mental event that had no clear temporal direction. For example, a control participant’s thought was, “I noticed this apparatus [EEG box] and I just imagined a picture in my mind in which that box was being used in a horror setting”. By contrast, a patient reported while noticing the same EEG box, “I wonder what this box with all these cables does. But I have no idea”. We display a detailed characterization of the temporal range of mind-wandering episodes in Figure 2. For statistical analysis, thoughts were binned into four main time categories, namely past (any thought related to earlier than the present moment), present (now), future (any thought related to later than the present moment), and atemporal thoughts.

**Figure 2.**
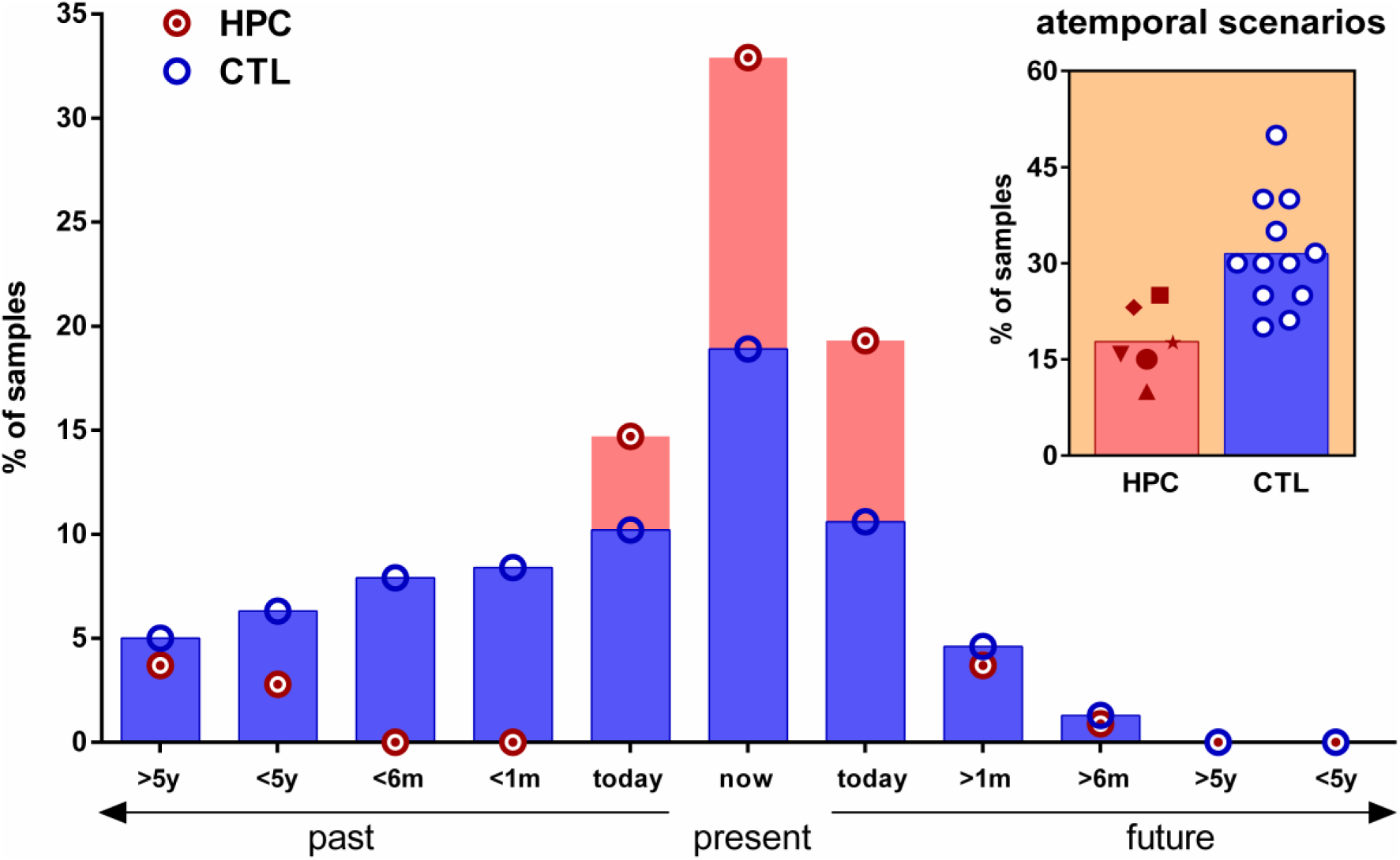
The temporal range of mind-wandering. Mean percentages of mind-wandering thoughts of patients with hippocampal damage (HPC, red circles with a dot) and controls (CTL, blue circles) for the past, present and future. For display purposes, thoughts are classified into time bins according to past (including earlier today), the present (now) and future (including later today); m=months, y=years. Control participants reported more thoughts related to the past than patients. In contrast, patients reported more thoughts related to the present than controls. The inset graph shows the percentage of thoughts during which patients (red symbols) and controls (blue circles) engaged in the imagining of atemporal scenarios.

#### Representation type

Thoughts were classified as either semantic or episodic (in line with established methods; (Levine et al., 2002; Andrews-Hanna et al., 2014b) and, in addition, whether they contained self-referential thinking or not (Andrews-Hanna et al., 2014b; Andrews-Hanna et al., 2014a). A thought was classified as semantic if it contained mentalization, or general knowledge about the world or the participant. For example, a semantic, self-referential thought of a patient was: “I am self-pondering. Am I a creative person?” A thought was classified as episodic if it contained specificity of time and place and a feeling of re- or pre-experiencing (Tulving, 1983, 2002). For example, an episodic, self-referential thought of a control participant was: “I am remembering a discussion I had with my friend at King’s Cross concourse a few weeks ago. I can see the scene clearly in front of me.” Of note, we also classified thoughts as episodic that had reference to a specific place and time, even if one or both were fictitious (time was more often fictitious). For example, an atemporal, episodic, non-self-referential thought of a control was: “I’m thinking about my friend. He’s travelling around giving lectures. I imagine an auditorium and see my friend speaking.”

#### Form of thoughts

We asked participants after each sample whether the thought had been verbal or visual, and if visual, whether it had been a scene or an object. Each thought was sorted into only one of these categories. Some participants reported that some of the visual scenes also contained verbal aspects, however, they regarded the visual scene as being more dominant. Therefore, these thoughts were classified as scenes. This classification was accomplished in agreement with each participant.

### Interrater reliability

In order to avoid potential rater biases, a second rater, who was blind to group membership, scored all thoughts from the patients and the control participants (except for one control dataset which was used as a training set). Interrater reliability was calculated as the direct correspondence between the two raters. That is, thoughts that were scored identically in a category were given a ‘1’ otherwise they were given a ‘0’. The reliability was then established as the sum divided by the total amount of rated thoughts. Therefore a value of 0.99 indicates that in 99% of samples the raters categorized them identically. The overall agreement between raters ranged between 84 and 99% across the thought categories (i.e., atemporal: 88%, coupled/decoupled 99%, semantic: 84%, episodic: 85%, and self-referential: 87%).

### Statistical analyses

Since most of the dependent variables did not meet the assumptions for parametric statistics, non-parametric tests were used for all within- and between-group analyses. Within-group analyses with more than two dependent variables were first conducted using Friedman tests (the non-parametric equivalent of repeated measures ANOVAs) and followed up with twotailed Wilcoxon Signed Rank tests (the non-parametric equivalent of paired t-tests). Between-group analyses with more than two dependent variables were first conducted using Kruskal-Wallis tests (the non-parametric equivalent of one-way ANOVAs) and followed up with twotailed Mann-Whitney U tests (the non-parametric equivalent of two-sample t-tests). Analyses with two dependent variables were directly compared using two-tailed Wilcoxon Signed Rank tests (within-group effects) or Mann-Whitney U tests (between-group effects). In all cases, we considered p-values less than 0.05 as statistically significant. For significant results we also report, where appropriate, the effect size (using non-parametric Cohen’s d) and we show the data of every participant.

## Results

### Frequency of mind-wandering

We first examined whether or not patients with hippocampal damage were able to mentally decouple from the current perceptual input (Fig. 1b, c). We found that the percentage of perceptually decoupled thoughts was greater than perceptually coupled thoughts in the controls (W=78.0, p=0.0005) and patients (W=21.0, p=0.03); Table 3). Notably, we found no difference between the two groups in the frequency of coupled (MWU=19.5, p=0.12) or decoupled (MWU=19.5, p=0.12) thoughts.

**Table 3.** Summary of mind-wandering data.

|  | CTL |  | HPC |  | p |
| --- | --- | --- | --- | --- | --- |
| **Mind-wandering** |  |  |  |  |  |
| *Perceptually coupled* | 6.8 | (7.5) | 13.4 | (13.6) | n.s. |
| *Perceptually decoupled* | 93.1 | (7.5) | 86.6 | (13.6) | n.s. |
| **Temporal range** |  |  |  |  |  |
| *Past* | 36.9 | (12.7) | 21.2 | (16.4) | * |
| *Present* | 18.9 | (8.3) | 32.9 | (7.4) | ** |
| *Future* | 15.6 | (9.9) | 24.6 | (9.2) | n.s. |
| *Atemporal* | 31.8 | (8.7) | 17.8 | (5.5) | **** |
| **Representational type** |  |  |  |  |  |
| *Episodic* | 72.6 | (12.4) | 24.8 | (14.3) | **** |
| *Semantic* | 27.4 | (12.4) | 75.2 | (14.3) | **** |
| *Self-referential* | 75.2 | (10.1) | 75.6 | (8.4) | n.s. |
| *Non-self-referential* | 27.8 | (10.1) | 24.4 | (8.5) | n.s. |
| **Form of thought** |  |  |  |  |  |
| *Scenes* | 63.5 | (5.4) | 12.3 | (11.7) | **** |
| *Objects* | 9.8 | (6.9) | 9.1 | (9.9) | n.s. |
| *Words* | 26.8 | (8.4) | 78.7 | (16.2) | **** |

For both groups, means (percentages) are displayed with standard deviations next to them in parentheses. CTL=healthy control participants; HPC=hippocampal-damaged patients; p=p-value for between-group non-parametric Mann-Whitney U tests. \*=p<0.05, \*\*=p<0.01, \*\*\*\*=p<0.001; n.s.=not significantly different.

### Temporal range of mind-wandering

Since mental time travel seems to occur frequently during mind-wandering (Smallwood and Schooler, 2015), we next examined whether the patient and control groups spontaneously thought about the past, present or future. After each thought sample, we asked participants whether the thought concerned the present moment, past or future, and if the latter two, how distant was it from the present moment (see Fig. 2 for a detailed visualization of multiple time bins). As described above, we also included an atemporal category in our analyses, comprising thoughts where a participant reported a mental event that had no clear temporal direction.

Examining the results for control participants in the first instance, we found that there was a significant effect of time category (Friedman statistic=19.99, df=3, p=0.0002). Post-hoc analyses showed that controls spent more of their mind-wandering time thinking about the past than the present (W=-62.0, p=0.002) or future (W=-72, p=0.002). They also spent more time simulating atemporal scenarios than thinking about the present (W=62, p=0.01). In contrast, there was no overall effect of time category for patients (Friedman statistic=6.86, df=3, p=0.07).

Direct comparison between the two groups revealed overall differences (Kruskal-Wallis statistic=31.93,df=7, p<0.0001). Post-hoc analyses showed that the patients thought less often than controls about past events (MWU=14.0, p=0.04, Cohen's d=1.1). By contrast, the patients thought more often about the present moment than control participants (MWU=7.0, p=0.0034, Cohen's d=1.7). There was no difference between the groups in the percentage of future-thinking, which was generally low for both groups (MWU=18.0, p=0.09). Lastly, we found that controls more often than the patients imagined atemporal events and hypothetical scenarios that concerned a fictitious reality, which was not attached to any temporal dimension (see the inset of Fig. 2, MWU=3.0, p=0.0007, Cohen's d=2.1).

### Representation type

We next investigated what the patients mind-wandered about (Fig. 3; Table 3). Focusing first on the control participants, we found that they reported significantly more episodic than semantic thoughts (W=78.0, p=0.0005), and more self-related than non-self-related thoughts (W=78, p=0.0005). The patients with hippocampal damage, on the other hand, experienced more semantic than episodic thoughts (W=-20.0, p=0.04), and more self-related than non-self-related thoughts (W=21.0, p=0.03).

**Figure 3.**
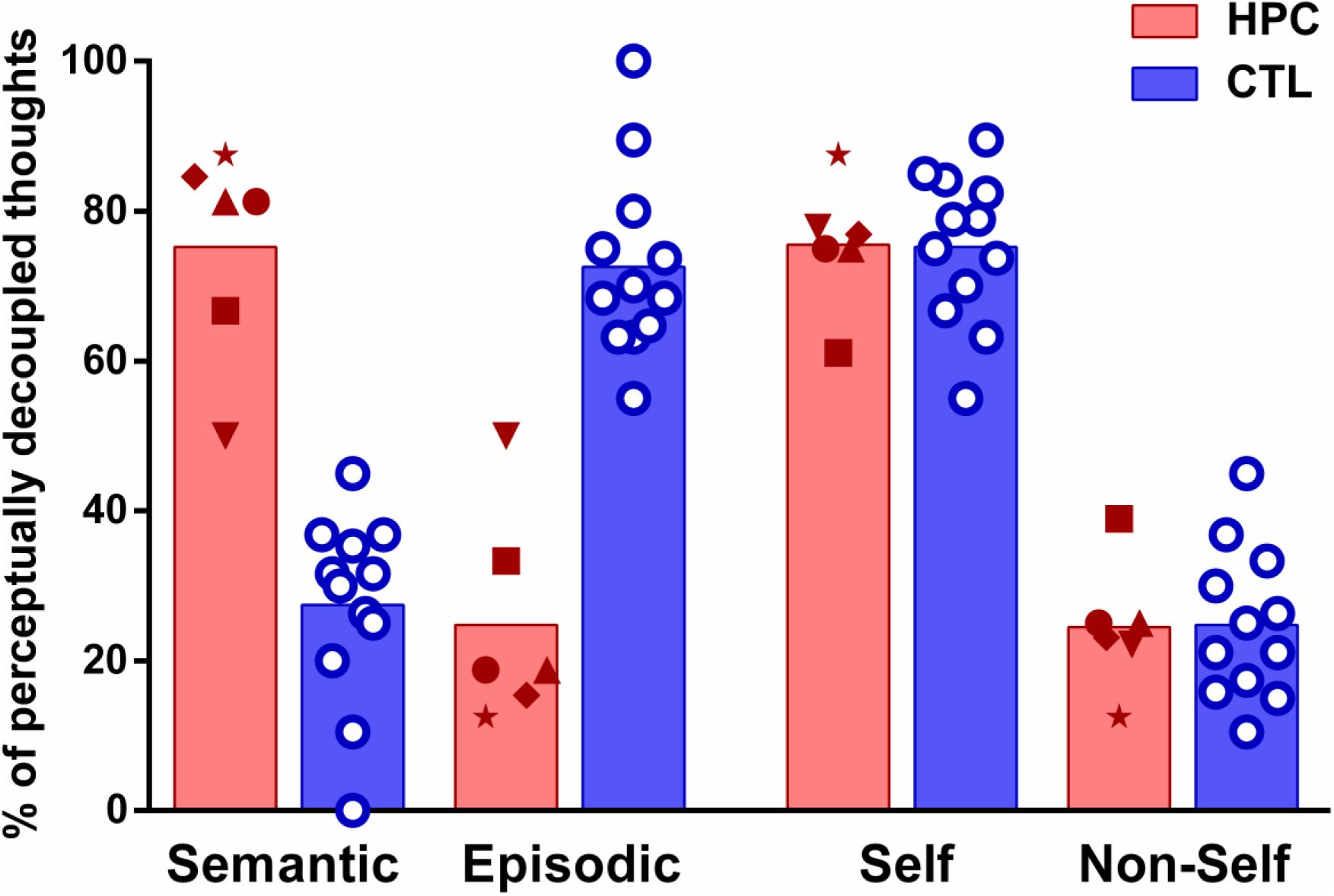
Semantic and episodic thinking during mind-wandering. Percentages of mind-wandering samples classified as semantic, episodic, self-referential or non-self-referential for patients with hippocampal damage (HPC, red symbols) and controls (CTL, blue circles). The patients had predominantly semantic thoughts, whereas the thoughts of the control participants were mainly episodic.

Directly comparing the participant groups revealed that the controls reported more episodic thoughts than the patients (MWU=0.0, p=0.0001, Cohen's d=2.6) and the patients reported more semantic thoughts than the controls (MWU=0.0, p=0.0001, Cohen's d=2.6). As expected, there was no significant difference in the percentage of self-referential (MWU=35.0, p=0.95) or non-self-referential (MWU=35.0, p=0.95) thinking between the groups. Together, these results show striking differences in the representational nature of spontaneous inner experiences between control participants and hippocampal-damaged patients.

### Form of thoughts

Finally, after each sample we asked participants whether the thought had been verbal or visual, and if visual, whether it had been a scene or an object (see Fig. 4, Table 3). For controls, we found overall differences in the frequency of the different forms of thought (Friedman statistic=21.83, df=2, p<0.0001). Post-hoc analyses showed that control participants reported that the majority of their thoughts involved visual scenes, more so than visual objects (W=-78.0, p=0.0005) or verbal thoughts (W=-78.0, p=0.0005), but more verbal thoughts than visual objects (W=58.0, p=0.007). For the patients too there were overall differences in the frequency of the different forms of thought (Friedman statistic=9.48, df=2, p=0.005). In striking contrast to controls, patients thought almost entirely verbally. They reported more verbal thoughts than visual scenes (W=21, p=0.03) and visual objects (W=21, p=0.03), with no difference between visual scenes and objects (W=-3, p=0.81).

**Figure 4.**
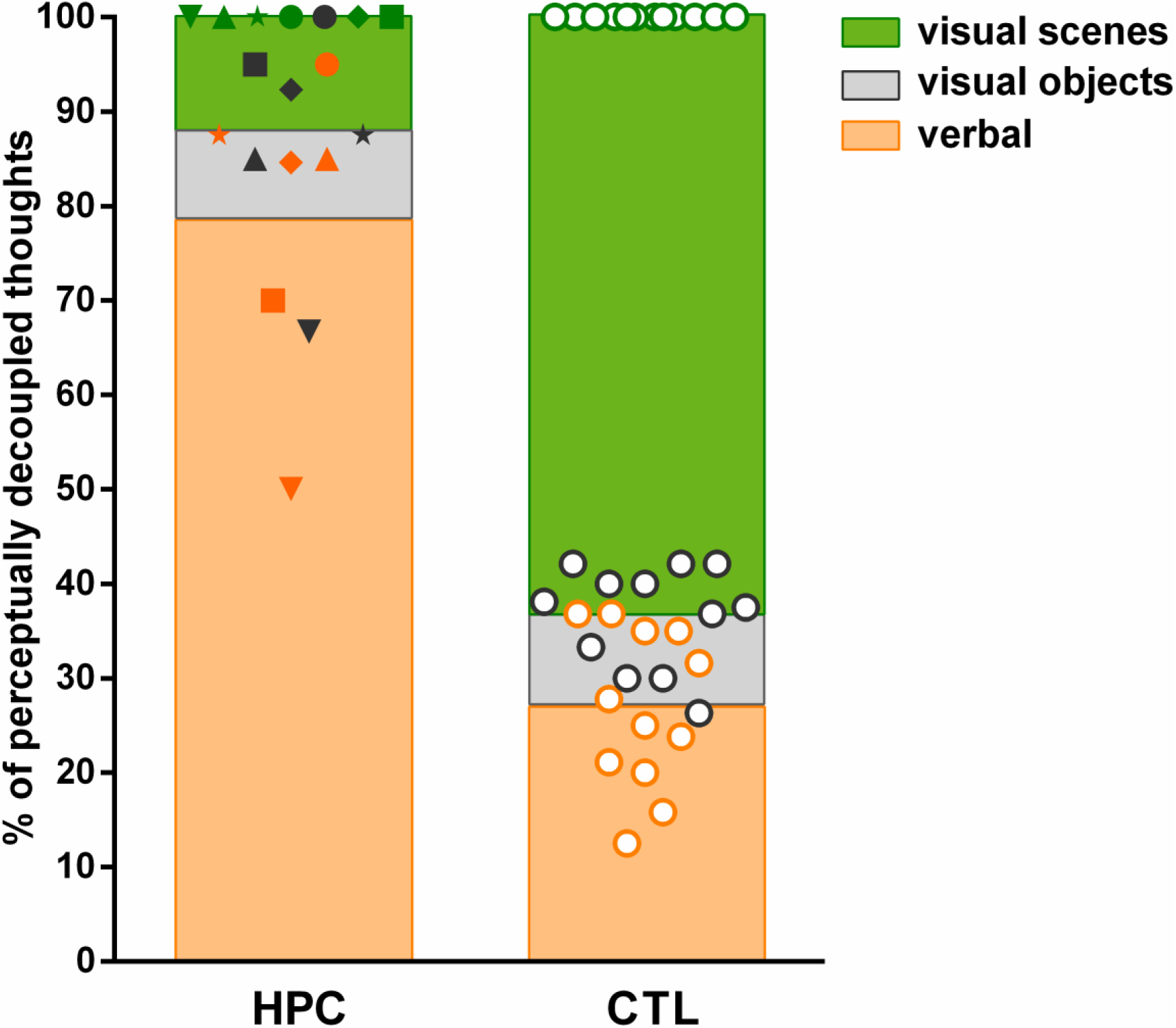
Cumulative percentages of visual and verbal mind-wandering thoughts. The average percentage of verbal thoughts is depicted per group (HPC=hippocampal-damaged patients, CTL=controls) as an orange bar; the individual data points are illustrated with orange symbols. The average cumulative percentage of thoughts containing visual objects is depicted as a grey bar above the average percentage of the verbal thoughts. The individual data points of thoughts containing visual objects (grey symbols) are illustrated as cumulative percentages above the orange data points (i.e., the patient represented as a square symbol reported around 70% verbal and around 25% visual object thoughts). Lastly, the average cumulative percentage of thoughts containing visual scenes is depicted as a green bar on top of the grey bar (green symbols all adding up to 100%). Whereas patients with hippocampal damage reported thinking in words for the majority of samples, healthy control participants’ thoughts were predominantly in the form of visual scenes.

These differences in the experiential form of mind-wandering were confirmed by directly comparing the participant groups (Kruskal-Wallis statistic=56.33, df=7, p0.0001). Whereas controls reported more visual scenes than patients (MWU=0.0, p=0.0001, Cohen's d=2.6), patients reported more verbal thoughts than controls (MWU=0.0, p=0.0001, Cohen's d=2.6), with no difference between participant groups for visual objects (MWU=30.0, p=0.59).

## Discussion

Mind-wandering is pervasive in humans and likely has an important role to play across cognition, influencing processes such as future planning, creative thinking and problemsolving (Baird et al., 2011; Baird et al., 2012; Andrews-Hanna et al., 2013). Here we showed that patients with hippocampal damage were able to perceptually decouple from the external world and experience spontaneous thoughts. Nevertheless, the small, selective lesions of their hippocampi dramatically affected the nature of their mind-wandering. Whereas healthy participants thought about the past, present and future, primarily in terms of episodic, detail-rich visual scenes, the patients mainly experienced verbally-mediated semantic thoughts anchored in the present. Previous studies have examined episodic thought processes in patients with hippocampal damage using explicit tasks, such as the Autobiographical Interview (Levine et al., 2002) or the scene construction task (Hassabis et al., 2007), that were designed to challenge the patients’ ability. In contrast, our findings show that even when there is no direct cognitive demand, the thought structure of people with hippocampal damage is strikingly different from healthy controls.

We first consider whether our results can be explained by a memory deficit that caused the patients to rapidly forget their mind-wandering thoughts before they could be accurately reported. We do not think is the case for a number of reasons. First, the patients had intact working memory and could retain task instructions during neuropsychological tests (Table 2) over longer time-scales than those in the current study. Second, we asked participants to describe two experiments unrelated to the current study shortly after completion, thus mirroring the timescale of reporting their mind-wandering experiences. All participants, including the patients, were able to provide accurate accounts of those experiments. Third, in previously-published studies involving the same patients and control participants using different paradigms, the patients were able to maintain information over time periods that were longer than those required for generating the current mind-wandering samples (McCormick et al., 2016, 2017a). Fourth, our sampling method did not involve any delay or distraction that might have affected the patients, nor did our protocol allow for increased post-hoc elaboration on the part of the control participants. Finally, if patients did not remember what they had been thinking about, the frequency of their mind-wandering would have been lower and they would have reported more mind-blanking, which was not the case. Thus, we are confident the patients were able to accurately report what was on their mind within seconds of the sampling cue.

Previous reports have estimated that humans tend to mind-wander about 30-50% of waking time (Kane et al., 2007; Killingsworth and Gilbert, 2010). Here, we report percentages nearer 80-90%. However, we specifically aimed to catch restful periods and so our higher percentage of mind-wandering thoughts suggests that we were successful at probing time points when mind-wandering levels were high.

Numerous studies have focused on delineating different aspects of inner experiences. For example, self-generated thinking (either intentional or unintentional; Seli et al., 2016) typically refers to the ability to mentally decouple from the current perceptual surroundings and generate independent internal thoughts (Smallwood and Schooler, 2015), which is a dichotomous definition that we employed in the current study. In reality, these self-generated thoughts align on a continuum ranging from closely task-related to totally task-unrelated (Smallwood and Schooler, 2015). What was most important for our research question was whether patients could decouple perceptually from their immediate surroundings in a completely task-free context. We found that they were able to do so and that the frequency of their mind-wandering did not differ from that of the control group. This result is especially noteworthy, given a recent study that found reduced frequency of mind-wandering in patients with ventromedial prefrontal cortex (vmPFC) lesions (Bertossi and Ciaramelli, 2016), a brain region with dense functional and anatomical connections with the hippocampus (Andrews-Hanna et al., 2010; Catani et al., 2012; Catani et al., 2013; McCormick et al., 2017b). Although there were differences in the experimental setup between our study and that involving the vmPFC patients, the difference in mind-wandering frequency observed in these two studies might indicate that the vmPFC is critical for the initiation of endogenous spontaneous thought and the hippocampus for its form and content.

At first glance, our finding of group differences in the temporal extent of mind-wandering is not surprising given the difficulty patients with hippocampal damage are known to have with recalling recent and remote episodic memories and imagining the future (Rosenbaum et al., 2008; Kurczek et al., 2015). However, these previous results were based on active and cognitively demanding tasks. To the best of our knowledge, this is the first indication that hippocampal-damaged patients experience reduced mental time travel even in their spontaneous thoughts. Of note, we did not replicate previous reports suggesting a near future-thinking bias in the mind-wandering of healthy participants (Stawarczyk et al., 2011; Song and Wang, 2012; Bertossi and Ciaramelli, 2016). The current experimental procedure and the older age of our participants may have influenced these results (Maillet and Schacter, 2016). For example, instead of sampling during low-demanding computer tasks or in natural environments that may encourage thoughts about the near future (e.g., “Where am I going after I’m finished here?”), we sampled thoughts across a structured day of stimulating research activities. This may have provided more opportunities to think about the recently-completed cognitive tasks or MRI scans. In addition, many previous studies have not included an atemporal category of thoughts, and it has been argued that thoughts labelled as future-oriented might in some instances be more accurately characterized as atemporal (Jackson et al., 2013). Indeed, in line with our results, it has been reported that healthy older adults experience more atemporal than future-oriented mind-wandering episodes (Jackson et al., 2013).

Recently, there have been increased efforts to map the complex cognitive processes that support mind-wandering to specific brain regions. While it is has been established that the DMN is associated with mind-wandering (Buckner et al., 2008; Andrews-Hanna et al., 2014a; Smallwood and Schooler, 2015), the contributions of specific brain areas within the DMN to mind-wandering remain unclear. Our results provide novel evidence that the hippocampus plays a causal role in episodic mind-wandering. These findings align with recent neuroimaging work that focused on a subsystem of the DMN, of which the hippocampus (and vmPFC) are nodes (Andrews-Hanna et al., 2010), and illustrated that functional and structural connectivity is stronger in individuals who report many detail-rich mental time travel experiences during mind-wandering (Karapanagiotidis et al., 2016; Smallwood et al., 2016). Our results further accord with network analyses in patients with hippocampal damage that showed altered hippocampal-neocortical connectivity patterns (Hayes et al., 2012; McCormick et al., 2014; Henson et al., 2016), which were associated with worse episodic memory capacity (McCormick et al., 2014). Of note, to the best of our knowledge, ours is the first report of a concomitant increase in spontaneous semantic thoughts associated with hippocampal damage. This may help to explain previous findings of increased connectivity between brain areas involved in semantic processing in resting-state fMRI studies involving similar patients (Hayes et al., 2012; McCormick et al., 2014).

In line with previous studies, our results demonstrate that mind-wandering episodes of control participants typically comprise visual imagery (Andrews-Hanna et al., 2013). We expand on existing studies by showing that visual imagery in task-unrelated mind-wandering of healthy controls primarily consists of spatially coherent visual scenes. In striking contrast, the patients with bilateral hippocampal damage no longer reported visualizing mental scenes, relying instead on a verbal thought structure. A scene construction deficit has been implicated in the impaired autobiographical memory and future thinking of patients with hippocampal damage (Hassabis and Maguire, 2007; Maguire and Mullally, 2013; Clark and Maguire, 2016). Our findings support this link between episodic thought and scene imagery. Importantly, this deficit also extends to scene perception tasks (Lee et al., 2005; Aly et al., 2013; McCormick et al., 2017a), suggesting that the lack of mental scenes is not because of faster visual degradation of imagery (Warren et al., 2011), but rather is due to an online scene construction problem. Thus, our results strongly suggest that hippocampal-supported scene construction is also central to the content and form of mind-wandering, and that without it, spontaneous thought seems to be reliant on verbal semantics.

Although the precise definition of mind-wandering is still debated, our results show that selective bilateral lesions to the hippocampus impair perceptually-decoupled inner thoughts in specific ways, thus informing the nature of mind-wandering and how it is realized at the neural level. That individuals with hippocampal damage experience mind-wandering but very little detail-rich mental imagery are important new insights which indicate the hippocampus is not necessary for the instigation of spontaneous thought per se. Instead, it seems to be crucial for processing the form and content of mind-wandering. Our results also speak to the functions of the hippocampus. By showing it plays a causal role in a phenomenon as ubiquitous as mind-wandering, this exposes the impact of the hippocampus beyond its traditionally-perceived role in memory, placing it at the center of our everyday mental experiences.

## Acknowledgments

We thank all the participants, particularly the patients and their relatives, for the time and effort they contributed to this study. We also thank the consultant neurologists: Drs. M.J. Johnson, S.R. Irani, S. Jacobs and P. Maddison. We are grateful to Martina F. Callaghan for help with MRI sequence design, Trevor Chong for second scoring the Autobiographical Interview, Alice Liefgreen for second scoring the mind-wandering thoughts, and Elaine Williams for advice on hippocampal segmentation. E.A.M. and C.M. are supported by a Wellcome Principal Research Fellowship to E.A.M. (101759/Z/13/Z) and the Centre by a Centre Award from Wellcome (203147/Z/16/Z). C.R.R. was supported by the National Institute for Health Research and John Fell OUP Fund. T.D.M. was supported by the Guarantors of Brain, the Patrick Berthould Charitable Trust and the Encephalitis Society.

